# Developmental Validation of the Illumina Infinium Assay using the Global Screening Array (GSA) on the iScan System for use in Forensic Laboratories

**DOI:** 10.1101/2022.10.10.511614

**Authors:** David A. Russell, Erin M. Gorden, Michelle A. Peck, Christina M. Neal, Mary C. Heaton, Jessica Bouchet, Alexander F. Koeppel, Elayna Ciuzio, Stephen D. Turner, Carmen R. Reedy

## Abstract

Microarray processing, which interrogates hundreds of thousands of single nucleotide polymorphisms (SNPs) across the human genome, has recently gained traction in forensics due to its use in forensic genetic genealogy, which is based on analysis using SNPs to compare distant relatives in publicly curated databases for the purposes of developing investigative leads or identifying human remains. To date, there has been no published developmental validation of microarray processing using the Scientific Working Group on DNA Analysis Methods (SWGDAM) Validation Guidelines for DNA Analysis Methods and Federal Bureau of Investigation Quality Assurance Standards. Validation of these methods are warranted to identify samples suitable for microarray analysis and to assess the quality of the data obtained prior to upload to genealogical databases. In this study, we validated the Global Screening Array (GSA) for use in forensic investigations according to SWGDAM guidelines, including the following studies: precision and accuracy, sensitivity, contamination, degradation, species specificity, mock case-type samples, mixtures, repeatability and reproducibility, and stability. Results indicated accurate genotype calls with SNP call rates >95% at DNA input as low as 0.20 ng. In addition to SNP call rate, we developed interpretation thresholds for signal intensity and heterozygosity to allow for sample quality assessment and identification of highly degraded and/or non-human DNA. This study demonstrates that high quality data can be generated from multiple sample types, including mock forensic evidence that simulated the challenges that are often encountered in forensic cold cases.

## Introduction

DNA profiling with short tandem repeats (STRs) continues to be the workhorse and gold standard in DNA forensics. STRs have been vetted through the court system and are widely accepted for DNA identification,^1,2^ providing high discriminatory value for source attribution/identification. However, due to the high mutation rate of STRs, kinship analysis is limited to close familial matches. Additionally, traditional STR profiles are uploaded to CODIS, a forensic database comprised of STR profiles from forensic evidence and known perpetrators, limiting the generation of new investigative leads to only those that hit to previously investigated crimes.

While STR typing remains the established method for human identification, these restrictions have resulted in a transition in the forensics community to single nucleotide polymorphism (SNP) genotyping, specifically for the generation of phenotypic trait assessment, biogeographical ancestry, and extended kinship comparisons. Most notably, SNP data generated from the DNA of suspects and missing persons has been uploaded to publicly available databases such as GEDmatch and used to identify distant relatives using forensic genetic genealogy (FGG).^3^ FGG gained attention in 2018 with the arrest of Joseph DeAngelo in the Golden State Killer case.^4,5^ In this case and many others, FGG has produced leads in cases that had gone cold or where traditional investigative means had been exhausted.^4,5^

The shift to SNP genotyping and FGG has largely been facilitated by the introduction of next generation sequencing (NGS) instruments and technology, including both microarray and sequencing workflows. The most popular method for generating SNP profiles has been using SNP microarray genotyping methods. Microarray technology generates data on hundreds of thousands of SNPs in a high-throughput, low-cost format. These high-density SNP profiles are designed to allow performance of distant kinship matching using publicly available databases and are the method most often used by the direct to consumer (DTC) genetic testing companies (e.g., Ancestry, 23andMe).

However, as microarray technology for use with FGG is often considered only for investigative lead generation with confirmatory testing performed using traditional STR typing, the forensic community has yet to establish standards and best practices for its use. Studies have begun to characterize the use of microarray systems with challenging forensic samples,^6,7^ but to date no developmental validation has been published by a forensic laboratory for application of microarray-based genome-wide SNP genotyping to forensic casework. Additionally, there exists minimal guidance, policy, or accreditation criteria for utilizing data generated from microarray workflows. The present study reports a developmental validation of the Infinium Global Screening Array (GSA; Illumina, San Diego, CA) using forensic validation guidelines and quality control measures. The GSA genotypes approximately 650,000 SNPs across the human genome and has been adopted globally in the fields of clinical disease research and consumer genomics. So much as it was applicable, the GSA developmental validation design was guided by the current Federal Bureau of Investigation (FBI) Quality Assurance Standards (QAS) for Forensic DNA Testing Laboratories and the Scientific Working Group on DNA Analysis Methods (SWGDAM) Validation Guidelines for DNA Analysis Methods.^8,9^ Here, we report on the following: precision and accuracy, sensitivity, contamination, degradation, species specificity, mock case-type samples, mixtures, repeatability and reproducibility, and stability. This study provides metrics and thresholds for evaluation and interpretation of microarray data obtained from the GSA that may provide guidance to forensic laboratories analyzing SNP genotyping data.

## Materials and Methods

### Samples and Study Design

The following DNA samples were obtained from the NIGMS Human Genetic Cell Repository at the Coriell Institute for Medical Research: NA12878, NA24385, and NA24631. A list of the Coriell and corresponding NIST samples used in this study can be found in Table S1. All samples in this validation were quantified using the Investigator Quantiplex^®^ Pro kit (QIAGEN, Hilden, Germany) on the 7500 Real-Time PCR System (Applied Biosystems™, Rockville, MD).

The precision and accuracy study evaluated NA12878, NA24385, and NA24631 at 200 ng total DNA input in duplicate. For the sensitivity study, three replicates of NA12878, were diluted to achieve total inputs of 200, 40, 20, 8.0, 2.0, 1.0, and 0.2 ng. A checkerboard pattern of alternating known human genomic controls at 200 ng and negative controls (NCs) was used at the initial setup of the Infinium assay to assess contamination (Figure S1). Additional NCs and reagent blanks (RBs) from other studies in the validation were included in the contamination assessment. For the degradation study, two DNA samples (NA12878 and NA24631) were exposed to UV-C (254 nm) to induce degradation (see supplemental methods). Four dosages and a no dosage control were selected for testing at 0.2 ng total DNA input in duplicate. For the species specificity study, purchased genomic DNA (Zyagen, San Diego, CA) from five non-human species (mouse, E. coli, rhesus monkey, yeast, and dog) were tested at 200 ng total DNA input in duplicate.

Mock case-type samples from a range of sources and of varying quality were obtained as extracts from an external agency. Out of 23 samples, 12 were selected for testing based on sample type, remaining extract volume, and DNA concentration. For the mixture study, five mixed DNA sample sets were created using NA12878 and NA24631 at the following proportions: 1:0, 9:1, 3:1, 1:1, 1:3, 1:9, and 0:1. Mixtures were processed in duplicate at 200 ng total DNA input. To assess repeatability and reproducibility, a subset of 12 and 13 samples, respectively, from the various studies above were re-genotyped. Finally, five BeadChips, totaling 102 unique samples, were re-scanned at various lengths of time after their initial scanning/analysis and data variability was assessed to establish the shelf life of GSA BeadChips for the purpose of possibly re-scanning at later dates. Lengths of times tested ranged from one day to two years post-original scan date. During the time between scan dates, BeadChips were stored at room temperature in a manner to protect them from light.

### SNP Genotyping and Analysis

Samples were prepared for SNP genotyping using Illumina’s Infinium HTS assay reference guide and manual protocol on the Infinium Global Screening Array-24 BeadChip, v3 (illumina, California, USA). All BeadChips were scanned on the Illumina iScan System array scanner according to the manufacturer’s protocol. Beeline was used to convert raw image (.idat) files to genotype call (.gtc) files.^10^ The data were analyzed using GenomeStudio^®^ 2.0 software (Illumina). ^11^

Following genome-wide genotyping in GenomeStudio, .gtc files were converted to variant call format (.vcf) files using Illumina’s GTCtoVCF software^12^. Call rate and total intensity were evaluated as metrics to assist with sample data quality evaluation. Call rate was determined as the number of SNPs yielding a genotype divided by the total number of SNPs interrogated. Total intensity was derived using the sum of the signal intensities for the red and green channels in the iScan. Genotype profiles from the NIST Genome-in-a-Bottle (GIAB) samples from the National Center for Biotechnology Information (NCBI)^13^ were used to generate GSA truth data (see supplemental methods for the data handling description). This data set was used in the precision and accuracy study to assess concordance. All other statistical analysis and visualization steps were performed using R.^17^

## Results

Sample information and GSA metrics for all samples included in this study are listed in Table S2.

### Precision and Accuracy

Two 200 ng replicates from NA12878, NA24385, and NA24631 were genotyped and compared against the truth GIAB genotypes. Call rates for all samples were >99%, meeting the manufacturer’s recommended metrics (Table 1A). Concordance at called sites was >99.9% for all samples, establishing high accuracy of the genotypes obtained on the GSA. Genotypes were also precise as concordance rates were >99.9% between GSA replicates of the same sample for all three samples tested (Table 1B).

**Table 1.** (A) Accuracy statistics for each sample when compared to Genome in a Bottle (GIAB)/NIST genotypes. Two replicates of each sample were compared. (B) Precision statistics between sample replicates.

| Sample ID | Call Rate (%) | SNPs Called | Concordance at Called Sites (%) | Missing SNPs | Missing SNP Rate (%) |
| --- | --- | --- | --- | --- | --- |
| NA12878 | 99.30 | 593,783 | 99.94 | 388 | 0.07 |
|  | 99.30 |  |  | 370 | 0.06 |
| NA24385 | 99.94 | 548,279 | 99.93 | 305 | 0.06 |
|  | 99.94 |  |  | 293 | 0.05 |
| NA24631 | 99.94 | 556,487 | 99.95 | 260 | 0.05 |
|  | 99.93 |  |  | 311 | 0.06 |

**B**
| Sample ID | SNPs Called | Concordance at Called Sites (%) | Missing SNPs | Missing SNP Rate (%) |
| --- | --- | --- | --- | --- |
| NA12878 | 593,783 | >99.99 | 634 | 0.11 |
| NA24385 | 548,279 | >99.99 | 499 | 0.09 |
| NA24631 | 556,487 | >99.99 | 476 | 0.09 |

### Sensitivity

Using genomic DNA from sample NA12878, three replicates were diluted for inputs at 200, 40, 20, 8.0, 2.0, 1.0, and 0.2 ng. Call rate and concordance metrics were calculated for each individual replicate, then averaged across all replicates within an input. Call rates were >99% for DNA inputs as low as 1.0 ng and >97% for inputs of 0.2 ng (Table 2). Comparing the genotypes to the 200 ng sample, results were highly concordant with >99% concordance at all DNA input amounts. While the 0.2 ng samples did not exceed the manufacturer’s recommended call rate of >99%, the high concordance indicates the genotypes are accurate.

**Table 2.** Call rate and concordance statistics for NA12878 at each DNA input level, averaged across replicates. The total number of SNPs genotyped is 635,930. The missing SNPs columns show the number/percent of SNPs where genotypes were missing in either the 200 ng sample (considered the truth data for comparison) or the lower input sample, preventing a concordance comparison across the two samples. The missing SNPs columns show the number/percent of SNPs where genotypes were missing in either the 200 ng sample (considered the truth data for comparison) or the lower input sample, preventing a concordance comparison across the two samples.

| Input (ng) | Concordant SNPs | Discordant SNPs | Concordance at Called Sites (%) | Missing SNPs | Call rate (%) |
| --- | --- | --- | --- | --- | --- |
| 40 | 635112 | 2 | >99.99 | 816 | 99.93 |
| 20 | 635065 | 4 | >99.99 | 861 | 99.92 |
| 8 | 634421 | 26 | >99.99 | 1484 | 99.82 |
| 2 | 633999 | 34 | >99.99 | 1896 | 99.76 |
| 1 | 631370 | 222 | 99.97 | 4338 | 99.37 |
| 0.2 | 615564 | 2000 | 99.69 | 18366 | 97.16 |

### Contamination

Twelve NCs and three RBs were assessed. The call rates and total fluorescence intensity from the iScan were compared to positive controls (NA12878, NA24385, NA24631) to determine differences between the sample types. As expected, the average call rate for the positive controls (200 ng) was 99.6%, but the NCs and RBs yielded much lower call rates ranging from 58-64%, (Table 3). The NC and RB call rates were higher than expected but can be explained by the way data is generated on the iScan. In short, SNPs are called based on relative fluorescence, and so samples with no true signal have genotypes called based on background noise, leading to inflated call rates. To further assess whether call rates are a result of true genotypes or background noise, the total fluorescence intensity was assessed on an individual sample basis. A clear difference in the total intensity between the positive and negative controls was demonstrated, with positive control samples generating intensity scores averaging >40,000 while the average for RBs and NCs was 958 (Table 3). From this, an intensity threshold (IT) was established using the intensity values from the NCs and RBs to distinguish true signal from noise when assessing unknown samples. The baseline noise value was calculated at a signal intensity of 1,400 by calculating 3 standard deviations above the mean NC value. To be conservative the IT was set at an intensity of 4,200, three times the baseline noise (Figure 1).

**Table 3.**
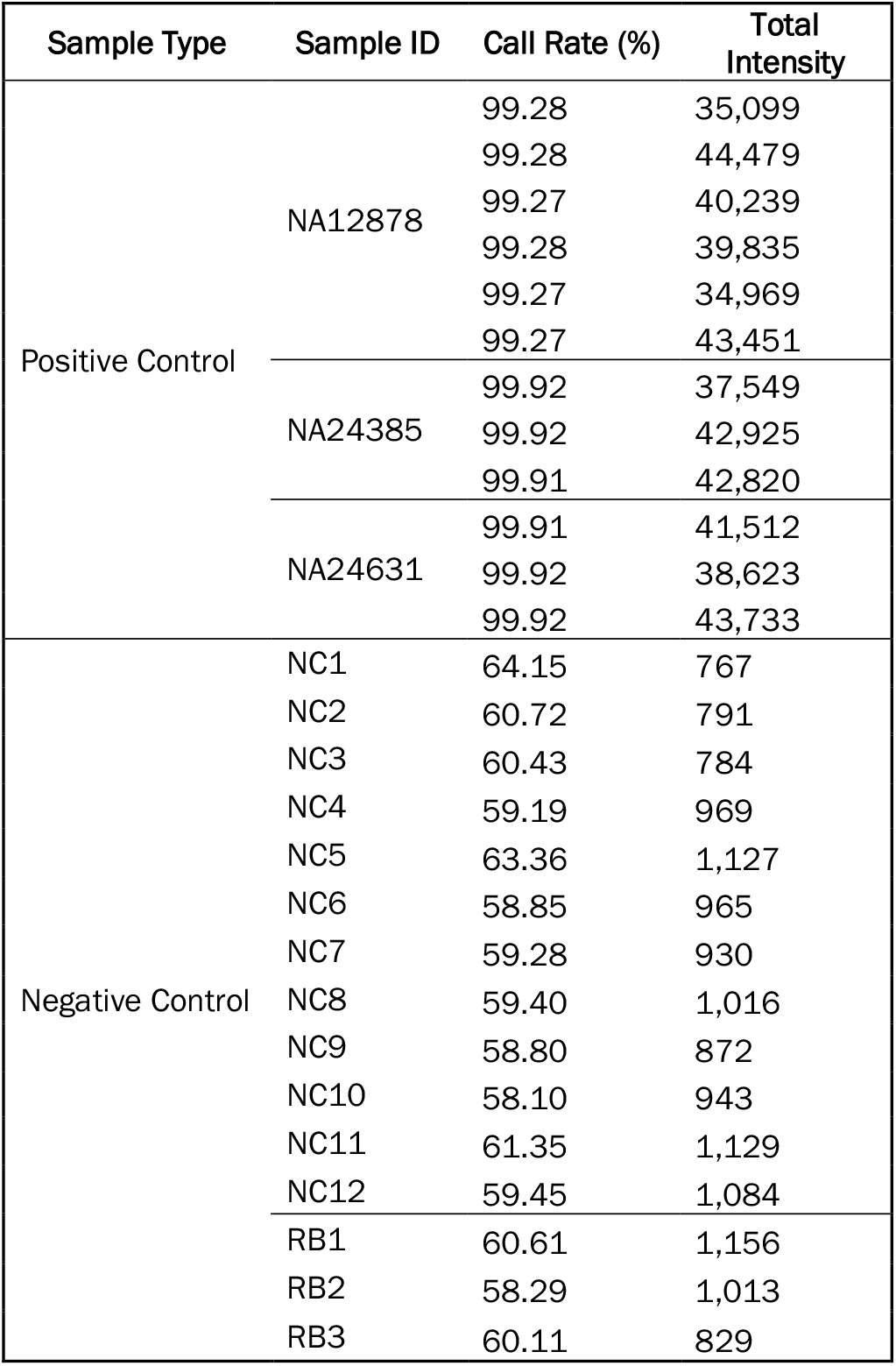
Call rates and total intensity for positive controls, negative controls (NCs), and reagent blanks (RBs). Positive control samples were run multiple times as part of various studies in this validation. The average call rate for negative controls was 60.3% (standard deviation (SD) = 1.87%), indicating a majority of genotypes are being called for these samples.

**Figure 1.**
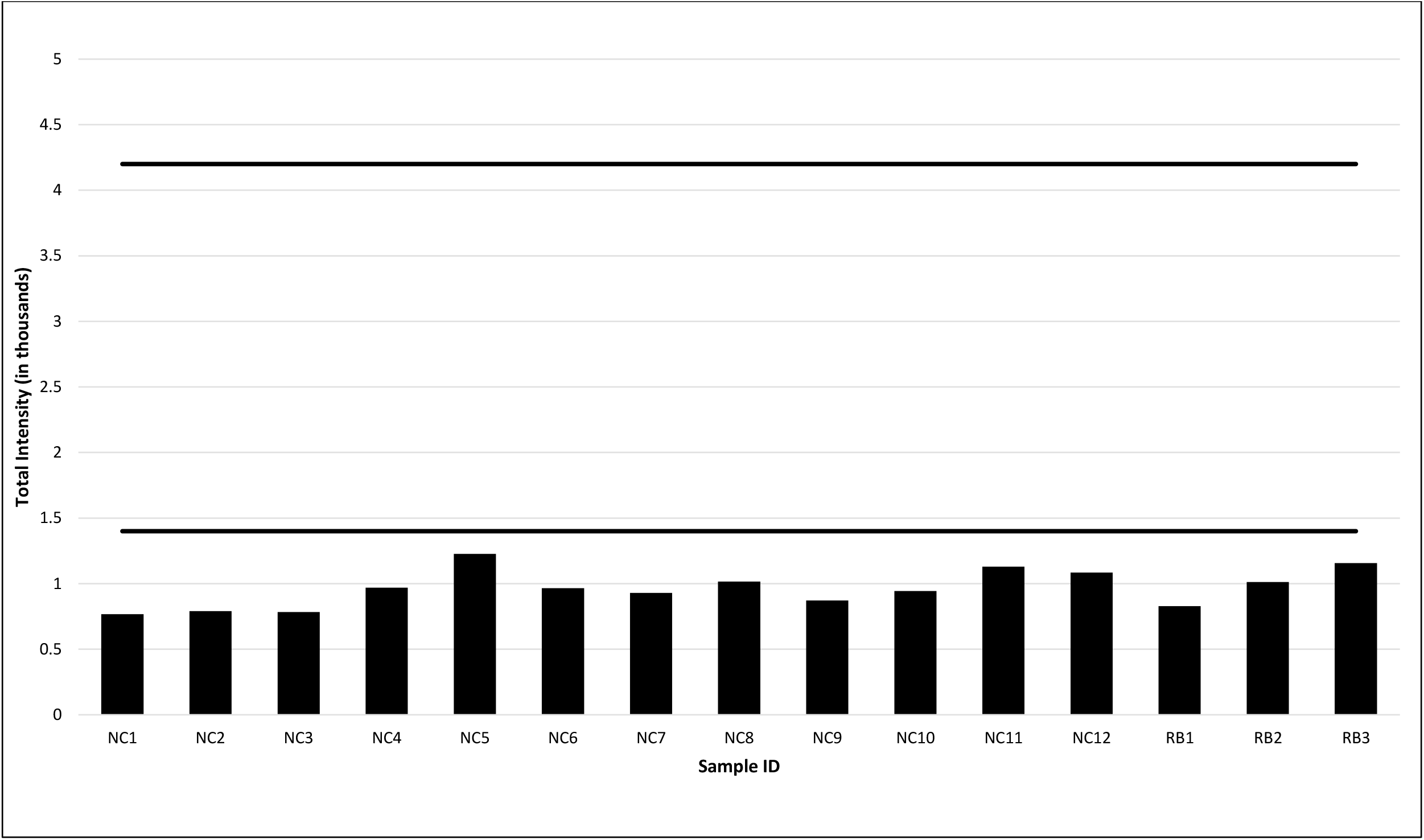
The mean total intensities for the 12 negative controls (NC) and three reagent blanks (RB) across all studies performed for this validation. The mean total intensity is calculated by summing the p95 red and p95 green intensity values from the GenomeStudio analysis. The baseline noise was calculated at 1,400 by taking 3 standard deviations above this mean (958) and rounding up to the next hundred. To be conservative, the intensity threshold was established at three times the baseline (4,200). Baseline and intensity thresholds are marked by solid black lines.

### Degradation

Samples were degraded using UV-C exposure at fixed dosages to assess sample quality at various degradation levels. Quantification of the pretreatment group was consistent with the targeted dilution of 0.05 ng/µL. Following UV-C treatment, quantification results for both samples showed an exponential increase in the degradation index (DI) (Figure 2A). The GlobalFiler data showed an increased ski-slope effect with increasing UV dosage, as expected (Figure 2B). The smaller amplicons exhibited moderate to high relative fluorescence units (RFU) throughout dye channels, then decreased in RFU with noticeable allelic drop-out in the larger amplicons. Dropout from DNA degradation increased proportional to UV-C exposure. Based on the observed degradation patterns in the electropherograms (data not shown), treatments 0, 125, 375, 625, and 1000 mJ/cm^2^ were selected for SNP genotyping.

**Figure 2.**
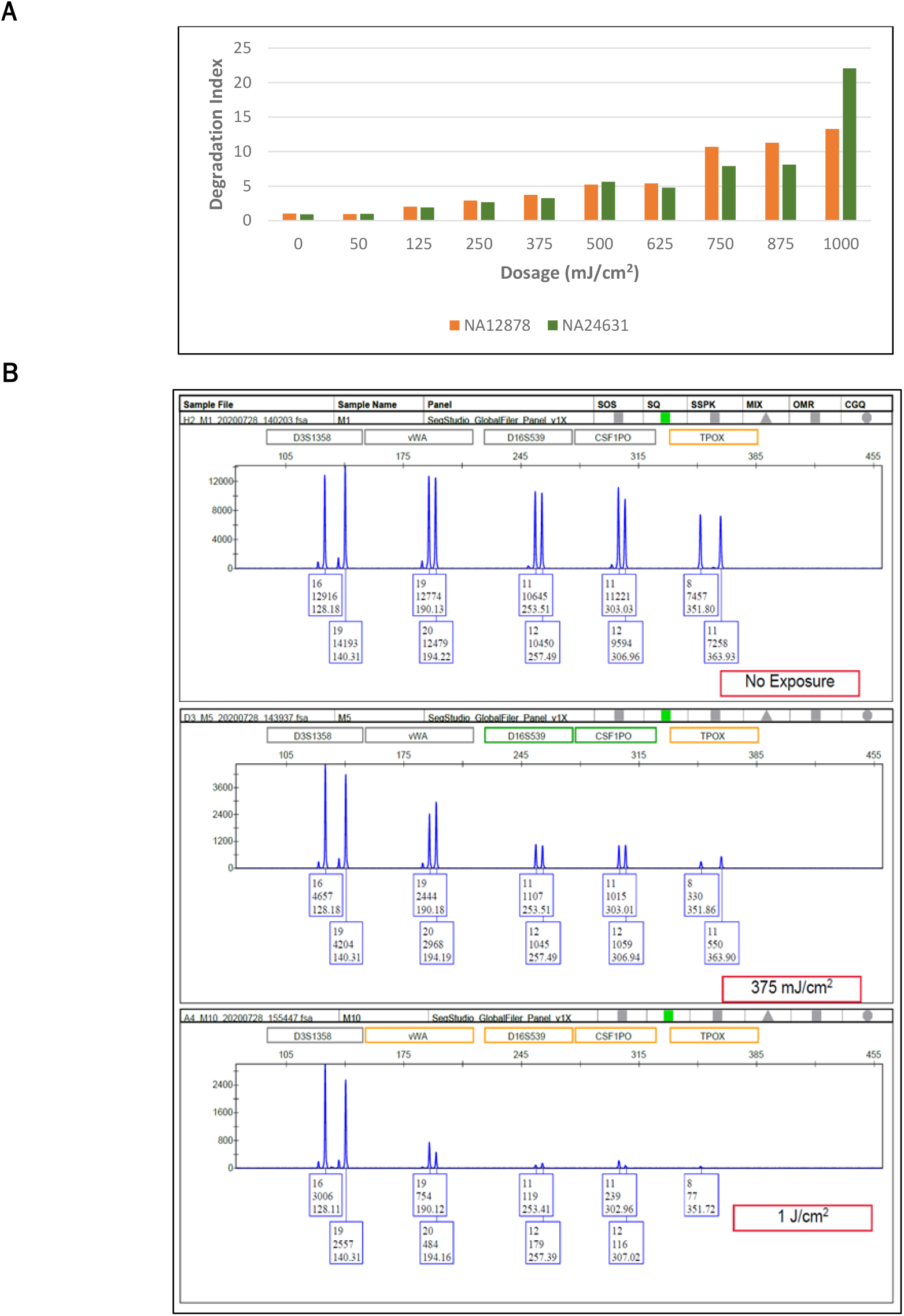
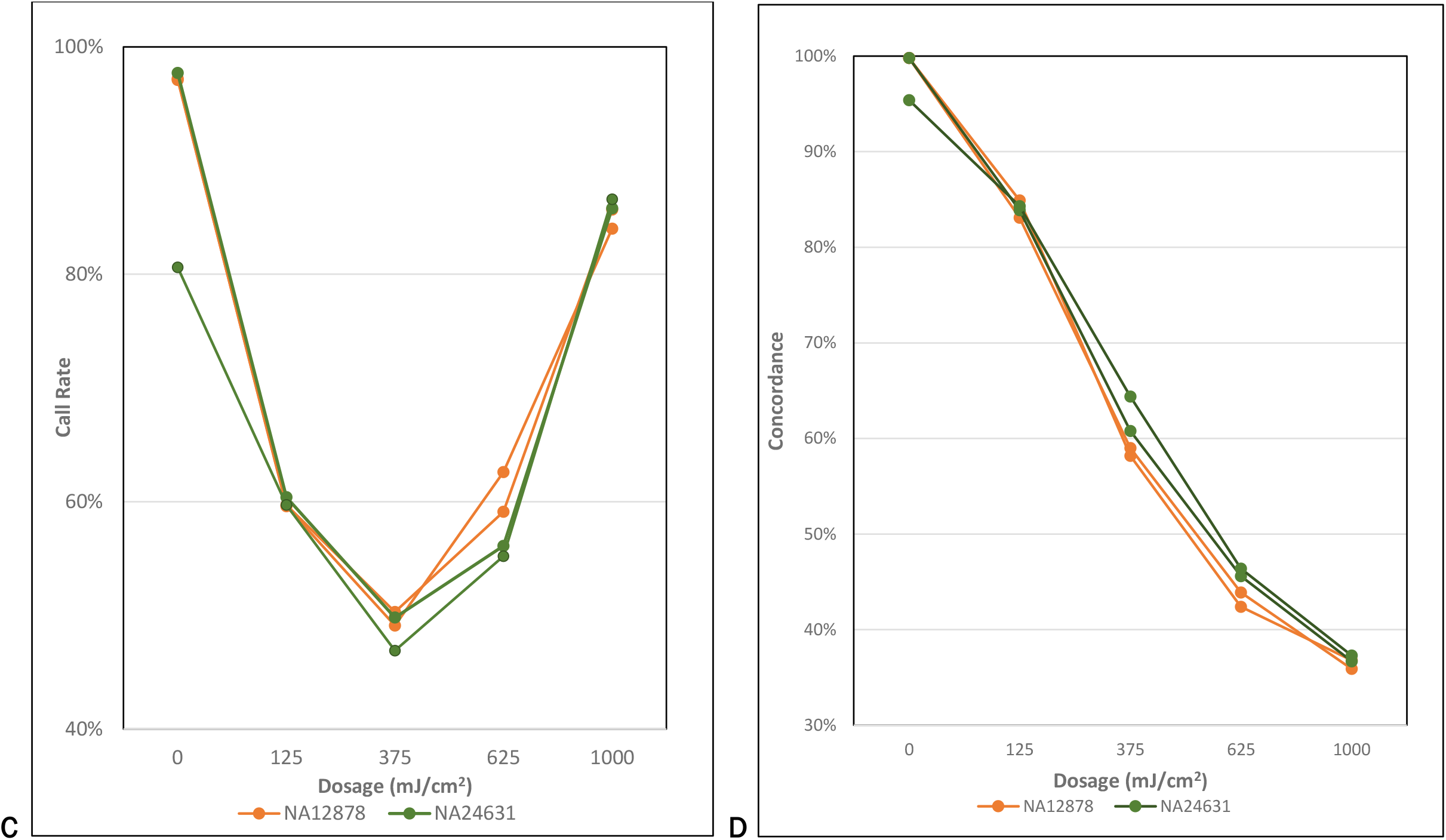
(A) Degradation index (DI) values across all dosage intervals. Dosage of 0 mJ/cm^2^ indicated control sample with no UV treatment applied. (B) Example of STR electropherograms showing increased degradation with increasing treatment dosages (0, 375, and 1000 mJ/cm^2^) in a single dye channel. The data was analyzed at a minimum analytical threshold of 50 RFU. “No exposure” demonstrates a ‘normal’ distribution of peak heights, while 375 mJ/cm^2^ treatment exhibits the ‘ski-slope’ pattern typically observed with degraded DNA samples. Treatment at 1000 mJ/cm^2^ exhibits the same sloped pattern in addition to allelic drop-out, suggesting greater DNA degradation. (C) Call Rates as it relates to the amount of UV dosage. Each line represents a replicate of the two DNA standards used in the study. (D) Sample concordance of each of the UV-C treated samples as compared back to the 20 ng control sample. Each line represents a replicate of the two DNA standards used in the study.

The SNP call rates in degraded samples initially decreased with the increase in the amount of dosage until approximately 375 mJ/cm^2^ and then began to rebound at higher dosages, resulting in similar call rates between the untreated (0 mJ/cm^2^) and the most degraded samples (1000 mJ/cm^2^) (Figure 2C). Yet, when concordance of the degraded samples was assessed against the untreated samples, concordance was less than 85% for all dosages (Figure 2D). Despite the high call rates, concordance decreased with increased degradation. This is most markedly observed in the most degraded extract of NA24631 where a call rate of 86.6% was observed but only 37% of calls were concordant with the untreated control.

The likely explanation for the increased call rate in highly degraded samples is allelic drop out, resulting in truly heterozygous loci genotyped as homozygous. To determine whether dropout was responsible for the observed results, sample heterozygosity was investigated. The rate of heterozygosity was calculated by dividing the number of autosomal heterozygous SNPs by the total number of called SNPs. The heterozygosity obtained for the unknown samples was then compared to the expected heterozygosity for GSA autosomal SNPs (excluding sex chromosomes and indels) in human populations from the 1000 Genomes data, consisting of 2,504 samples from 26 populations. The heterozygosity observed in the 1000 Genomes samples ranged from 15.2-19.4%, averaging 17.3 ± 0.684%. From these values a heterozygosity threshold range of 15-20% was set using three standard deviations above and below that mean.^14^ Samples falling outside this range were considered unreliable and not considered for downstream analysis. Degraded samples with dosages of ≥375 mJ/cm^2^ demonstrated heterozygosity levels <15% (Figure 3), indicating significant allelic dropout, and signifying that call rate alone is not sufficient for evaluating the accuracy of SNP microarray data.

**Figure 3.**
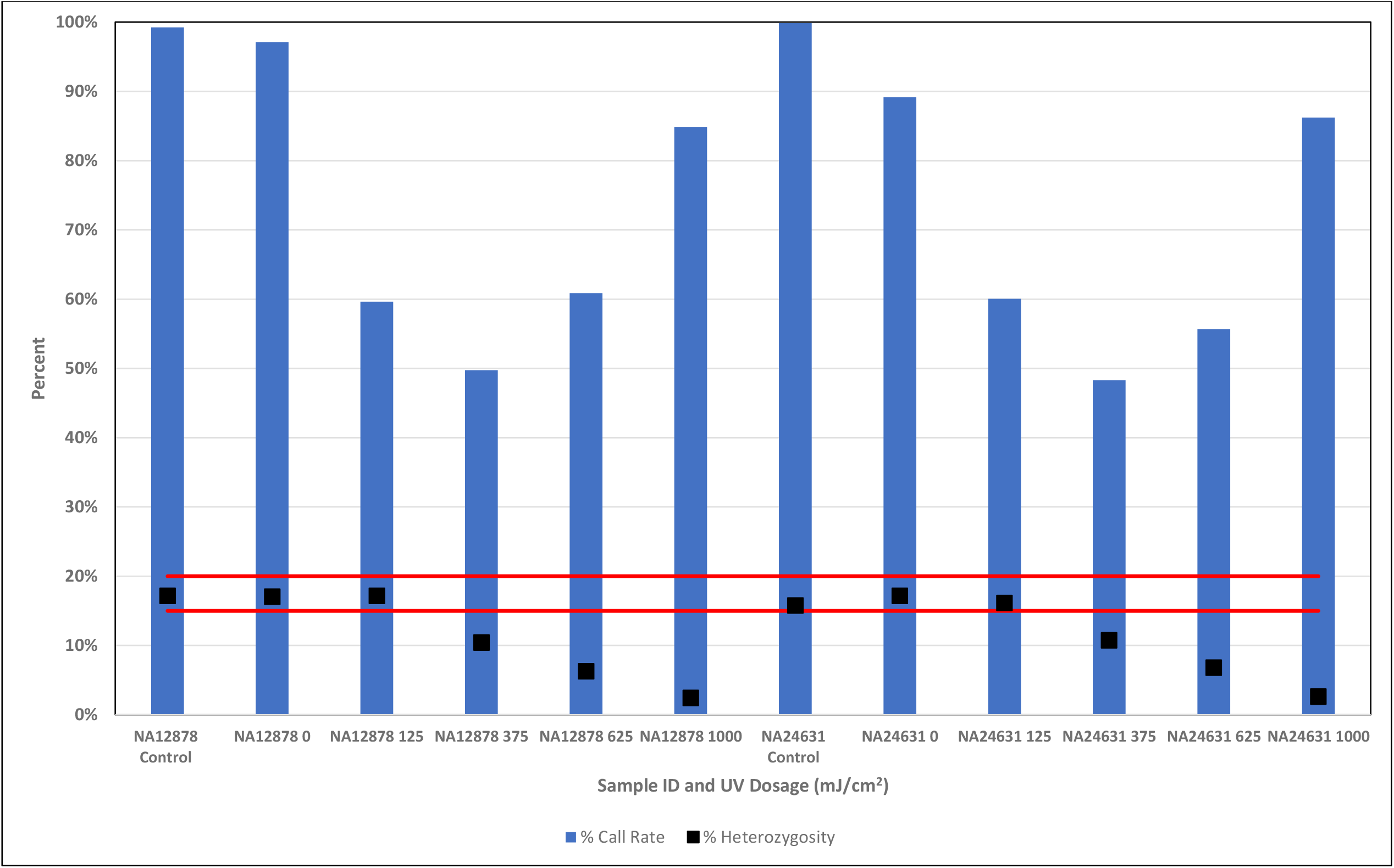
Heterozygosity and call rate of degraded samples, averaged across two replicates for each dosage. Sample IDs are listed as the amount of dosage for each (e.g., 125=125 mJ/cm^2^ dosage). All samples met the total intensity threshold with the exception of one replicate of NA12878 with 1000 mJ/cm^2^ of UV-C degradation. The red lines represent the heterozygosity upper and lower bounds.

As a result of this observation, heterozygosity was assessed on all previously analyzed samples and subsequent studies to evaluate its effectiveness as an indicator of sample quality. While most samples were in the targeted heterozygosity range (N=94), 42 fell outside the established heterozygosity threshold and were appropriately flagged as unreliable profiles. In addition to the degraded samples, flagged samples included NCs and RBs, non-human DNA samples, and low quantity (i.e., <0.2 ng) samples (Table S3).

### Species Specificity Study

A combination of intensity, call rate, and heterozygosity were used to distinguish between non-human and human samples tested in this study. While the call rates for the non-human samples were similar to those of RBs and NCs for most samples, the call rate was unexpectedly high for dog DNA (average of 81.5%). However, these samples yielded low intensity (2,194) and a heterozygosity significantly below the expected range (1.65%). In fact, the only species tested with a total intensity exceeding that of background noise was the rhesus monkey, due to potential sequence similarity with humans. However, call rate and heterozygosity statistics for rhesus monkey replicates averaged 62.6% and 5.6%, respectively. In sum, all non-human samples tested have either total intensity below threshold and/or heterozygosity outside of the range expected for authentic human samples. Consequently, these samples would not progress for upload to a genealogical database.

### Mock Case-Type Samples

DNA extracts used in the mock case sample study were obtained from an external agency and originated from various sources. All four RBs associated with these samples yielded no quantification results and thus were not genotyped. Of the 23 samples submitted, nine had sufficient concentration (>0.05 ng/μL) to meet the minimum total DNA input established from the sensitivity study (0.20 ng) and were selected for genotyping with the GSA. An additional three bone samples yielded concentrations slightly below the threshold (0.13 ng on average) and were also genotyped. The bone samples were included because bone was not represented in samples above the minimum input requirement, and it was of interest to assess the feasibility of performing SNP genotyping on this sample type. These samples also provided further data to assess DNA input into the GSA below 0.20 ng.

While DNA input into the GSA does seem to have some impact on call rate, there is no direct correlation (Table 4; Figure S2). Call rates ranged from ∼68-93% for samples that met the established DNA input of 0.2 ng, except for Mock13, which encountered a possible genotyping scan error on the iScan. All these samples also passed the IT threshold and were within the expected heterozygosity range. The three samples with a DNA input below 0.2 ng yielded call rates of 39-68%, and intensity values consistent with baseline noise. These results demonstrate that the validation protocol and interpretation thresholds developed distinguish between samples that have generated quality, uploadable data and those that have not.

**Table 4.**
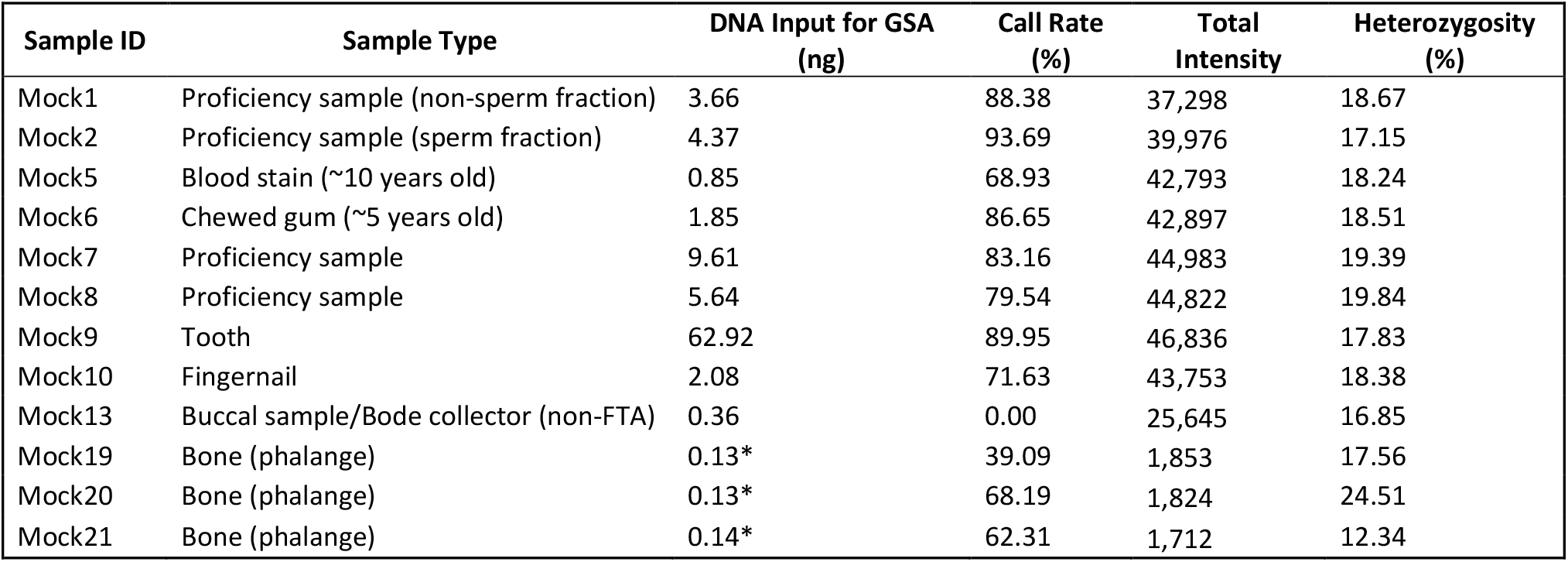
Sample information and GSA metrics for mock case samples. Any samples not meeting the 0.2 ng DNA input requirement are indicated with an asterik (*). Sample Mock13 generated a call rate of 0% despite a high total intensity, and likely represents a genotype processing error on the iScan that could be rectified with rescanning or reprocessing.

### Mixtures

Genomic DNA from two DNA standards of divergent ancestry (NA12878, NA24631) was prepared at different mixture ratios and genotyped using the GSA. The call rates from the mixture samples ranged from 81.8-99.9%, with the lowest call rates observed in the 1:1 mixtures (∼82%) and increasing call rates with increasing donation by the major contributor (Table 5). The lower call rate for 1:1 mixtures can be explained by the genotyping method used by the iScan and GenomeStudio. Genotypes are determined by the relative fluorescence of two dyes, which allow loci to be graphed on a plot that bins genotypes into one of three clusters (AA, AB, or BB). Calls that do not fall neatly into one of the expected clusters results in no call at that locus (e.g., Figure S3). Therefore, in a 1:1 mixture where one sample genotypes as AA and another as AB, for example, the ratio of A:B alleles is 3:1. GenomeStudio, which uses previously defined clusters based on single source samples, may incorrectly call this locus, or fail to call it at all.

**Table 5.** Results of the mixture study, including the mixture ratio and associated statistics for each of the mixture samples. Each sample was run in duplicate. Total DNA input was 200ng for each of the mixture samples. Concordance was calculated compared to the profile of the major donor. No concordance was calculated for the single source samples (NA12878 and NA24631). Concordance was assessed to both donors for 1:1 mixtures.

| Sample ID | Mixture Ratio (NA24631:NA12878) | Call Rate (%) | Heterozygosity (%) | Concordance to Major Donor (%) |
| --- | --- | --- | --- | --- |
| NA12878 | 0:1 | 99.29 | 17.21 |  |
|  |  | 99.30 | 17.21 |  |
| NA24631 | 1:0 | 99.94 | 15.89 |  |
|  |  | 99.94 | 15.88 |  |
| Mix1 | 3:1 | 87.55 | 19.12 | 97.92 |
|  |  | 87.66 | 19.11 | 97.85 |
| Mix2 | 9:1 | 97.20 | 16.21 | >99.99 |
|  |  | 97.16 | 16.20 | >99.99 |
| Mix3 | 1:1 | 81.82 | 16.46 | 90.19 ; 93.24 |
|  |  | 82.36 | 17.08 | 89.92 ; 92.90 |
| Mix4 | 1:3 | 90.25 | 18.85 | 99.80 |
|  |  | 90.21 | 18.79 | 99.72 |
| Mix5 | 1:9 | 98.45 | 17.33 | >99.99 |
|  |  | 98.50 | 17.32 | >99.99 |

Concordance of the mixed samples was calculated to the major donor genotypes (Table 5). Concordance with the major donor was >97% for all mixture ratios other than 1:1 mixtures. For the 1:1 mixture ratio the concordance to either donor fell below 95% ranging from ∼89-93%, indicating detection of the minor contributor. Mixtures of 3:1 yielded ∼97-99% concordance to the major donor, and 9:1 mixtures demonstrated 100% concordance with the major genotype. However, it is important to note that this may differ depending on DNA quality and quantity as well as allele sharing between the individual donors at GSA genotyped SNPs.

While the call rate, intensity, and heterozygosity metrics were not able to distinguish mixtures from single source samples (Table S2), this study has shown that in instances where the mixture ratio is 3:1 or greater, the produced genotype of the major contributor is accurate without requiring removal of the minor contributor. Therefore, case circumstances may exist where processing may move forward for mixed samples if the minor contributor is determined to be low and genotyping for the major contributor will not be impacted.

### Repeatability and Reproducibility

Samples from multiple studies, including positive controls and mock samples, were processed in replicate to demonstrate the repeatability and reproducibility of the assay. Metrics (i.e., call rate, intensity, and heterozygosity) were similar between multiple replicates of the same sample. Minor differences in these metrics between replicates can be attributed to scan variability and/or variability associated with low input and degraded samples. The concordance was calculated between replicates of the same sample, and on average reached >99.9% and >97% in pristine and mock casework samples, respectively (Table S4). Therefore, data generated from the GSA workflow has shown to be reproducible.

### Stability

Five BeadChips were re-scanned at lengths of time ranging from one day to two years after the original scan date (Table S5). No discernable difference was observed between the call rates and heterozygosity (Figures S4 (A) and (B), respectively) generated from the original scan and re-scan data for 124 samples used in this study. A large majority (n=105) of the samples tested (84.7%) differed by <1% in the call rate calculated between the two scans. Similarly, 117 of the samples tested (94.4%) differed by <1% in the heterozygosity calculated between the two scans. The remaining samples that displayed >1% difference in either heterozygosity or call rate represent degraded and non-human samples that would not have exceeded established thresholds (i.e., signal intensity, call rate, and heterozygosity) on the initial scan and thus would be unlikely candidates for re-scan. Overall, the results from this study showed minimal difference in GSA metrics when stored at room temperature for up to two years.

## Discussion

This developmental validation demonstrates the efficacy of the GSA to provide high quality SNP data from sample types common in forensic investigations. These studies show high precision and accuracy with DNA input quantities down to 0.2 ng, much lower than the manufacturer’s recommendations. The results presented herein demonstrate that call rate alone was not sufficient as the sole metric to evaluate data quality of forensic samples from the GSA. This validation resulted in the derivation of two additional metrics from the produced SNP data that have shown to give additional insight into data quality and can be used to determine sample eligibility into downstream databases. These metrics, combined with the call rate, help identify low quality data that may result from low quantities, degraded, or non-human DNA.

The contamination assessment flagged that call rate alone is insufficient as a metric, as the NCs and RBs gave higher call rates than expected (58-64%). The iScan system uses relative fluorescence signal to call genotypes, and BeadChip technology is not designed for empty wells. Thus, non-specific binding or background noise artificially inflates SNP call rate as the resolution algorithm tries to maximize signal detection. To account for this, the inherent baseline noise produced during imaging using NCs was used to set an intensity threshold at three times the baseline noise level. The intensity threshold was applied to all samples used in this study and was able to filter out severely degraded samples with sufficient DNA quantity, NCs, and most non-human samples (Table S2).

The heterozygosity threshold originated from the degradation study due to observed high call rates in highly degraded samples. Concordance data shows that the accuracy of the genotype calls in these highly degraded samples is low, indicating allelic dropout. Application of a heterozygosity threshold range, calculated for human populations on GSA-specific SNPs, allows for an assessment of possible degradation in unknown samples that could result in upload of incorrect genotypes to a genealogical database. This metric can be utilized to distinguish samples of non-human origin where call rate and intensity values may suggest otherwise (e.g., rhesus monkey).

Data produced from the GSA should be assessed using the above-stated metrics. Samples producing call rates lower than 60% are not reliable and should not be uploaded to FGG databases, while samples exceeding 60% call rate should undergo additional analysis. Our data driven, hierarchical, approach seeks to maximize the samples eligible for GSA SNP genotyping while minimizing the upload of poor-quality data to genealogical databases that can result in fortuitous matches, unnecessarily excessive genealogical research, and other detrimental downstream effects.

## Conclusions

This validation was performed to establish use of the Illumina GSA for forensically relevant sample types under the guidance of current FBI QAS for Forensic DNA Testing Laboratories^8^ and SWGDAM Validation Guidelines for DNA Analysis Methods^9^. The validation shows that high quality results can be obtained with a DNA input of 0.2 ng, significantly less than the manufacturer recommended input of 200 ng. Our validation demonstrates that call rate alone is not sufficient to assess data quality. Heterozygosity and total intensity metrics should be used in addition to call rate to assess sample quality prior to upload to genealogical databases.

## Author Contributions

**David A. Russell:** Conceptualization, Methodology, Software, Validation, Formal Analysis, Investigation, Resources, Data Curation, Writing -Original Draft, Writing - Review & Editing, Visualization, Supervision, Project Administration.

**Erin M. Gorden**: Methodology, Validation, Formal Analysis, Investigation, Data Curation, Writing - Original Draft, Writing - Review & Editing, Visualization.

**Michelle A. Peck**: Methodology, Validation, Formal Analysis, Investigation, Data Curation, Writing - Original Draft, Writing - Review & Editing.

**Carmen R. Reedy**: Conceptualization, Methodology, Writing - Original Draft, Writing - Review & Editing, Supervision, Project Administration.

**Stephen D. Turner**: Conceptualization, Methodology, Software, Validation, Formal Analysis, Investigation, Resources, Data Curation, Writing - Review & Editing.

**Christina M. Neal**: Conceptualization, Methodology, Writing - Review & Editing, Supervision.

**Mary C. Heaton**: Conceptualization, Methodology, Writing - Review & Editing, Supervision.

**Alexander F. Koeppel**: Writing - Review & Editing.

**Jessica Bouchet**: Methodology, Validation, Investigation. Elayna Ciuzio: Conceptualization, Methodology, Supervision.

