## Supplemental Information for "Developmental Validation of the Illumina Infinium Assay using the Global Screening Array (GSA) on the iScan System for use in Forensic Laboratories"

### SUPPLEMENTAL METHODS

#### Degradation

NA12878 and NA24631 were normalized to 0.05 ng/µL in a volume of 20 µL, of which 2 µL was used for pre-UV quantification. Once prepared, samples were covered in foil to initiate control treatment of UV exposure, prevent ambient light exposure, and were stored at 4°C.To perform UV-C (254 nm) degradation, the foil was removed from the samples, and the UV dosage applied at intervals of 50, 125, 250, 375, 500, 625, 750, 875, and 1000 mJ/cm$^2$. A dosage of 0 mJ/cm$^2$ acted as a control. Quant values were obtained for each aliquoted sample both pre- and post-UV treatment. Genomic DNA was amplified using the GlobalFiler™ (Applied Biosystems) PCR amplification kit and typed on the SeqStudio™ (Applied Biosystems). Genetic Analyzer data was analyzed using GeneMapper™ ID-X software v1.6 (Applied Biosystems). Treated samples that covered a range of degradation in the STR profile interpretation, along with the 0 mJ/cm$^2$ dosage controls, proceeded to microarray analysis in duplicate. An additional dilution was prepared from each of the DNA standards at a concentration of 20 ng/µL to act as a control and was compared to the treated samples to assess concordance.

#### GIAB Truth Data Set for Concordance

To assess accuracy, the NIST/Genome-in-a-Bottle (GIAB) gold standard genotypes were used to generate GSA truth data. The GIAB variant call format (VCF) files were first downloaded from NCBI. GIAB profiles are obtained through whole genome sequencing and therefore reported data only includes calls at sites which are variant with respect to the reference genome. To obtain genotypes at all GSA sites (including those that may be invariant), bcftools[1] was used to call homozygous reference genotypes at invariant positions when the sequenced region was of high-confidence. The NA12878, NA24385, and NA24631 replicates were compared to the truth sets for concordance using bcftools gtcheck.[1] Comparison of the generated profiles to the GIAB profiles excluded SNPs that did not genotype in any sample, that did not genotype in the NIST/GIAB sample, or that are not biallelic). Concordance statistics were calculated for each donor replicate, then averaged across all replicates. For the precision analysis, genotypes for the replicate samples were compared against each other for all SNPs used in the accuracy analysis.


### SUPPLEMENTAL TABLES & FIGURES

**Table S1**

Coriell and NIST sample IDs derived from the same individual.

| Sample ID | |
|---|---|
| Coriell | NIST |
| NA12878 | HG001 |
| NA24385 | HG002 |
| NA24631 | HG005 |

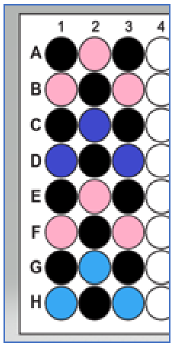


**Figure S1**

Contamination study plate layout and placement of the DNA positive and negative controls (NCs). Pink=NA12878; Purple=NA24385; Blue=NA24631; Black=NCs

**Table S2**

Sample information and GSA metrics for all samples assessed in this study. Repeatability samples were included as part of various other studies (e.g., replicates of NA12878 in the sensitivity study) and are therefore not listed here specifically as repeatability.

| Study | Sample ID | DNA Input for GSA (ng) | Call Rate (%) | Total Intensity | Heterozygosity (%) |
|---|---|---|---|---|---|
| Accuracy and Precision | NA12878 | 200 | 99.30 | 42,663 | 17.20 |
| | | 200 | 99.30 | 33,908 | 17.20 |
| | NA24385 | 200 | 99.94 | 31,771 | 16.80 |
| | | 200 | 99.94 | 32,834 | 16.80 |
| | NA24631 | 200 | 99.94 | 34,226 | 15.90 |
| | | 200 | 99.93 | 35,977 | 15.90 |
| Sensitivity | NA12878 | 200 | 99.29 | 28,611 | 17.20 |
| | | | 99.29 | 29,814 | 17.20 |
| | | | 99.28 | 30,826 | 17.20 |
| | | 40 | 99.28 | 32,142 | 17.20 |
| | | | 99.29 | 39,945 | 17.20 |
| | | | 99.30 | 35,823 | 17.20 |
| | | 20 | 99.29 | 33,741 | 17.20 |
| | | | 99.27 | 40,821 | 17.20 |
| | | | 99.29 | 37,013 | 17.20 |
| | | 8 | 99.25 | 36,112 | 17.20 |
| | | | 99.04 | 39,697 | 17.20 |
| | | | 99.26 | 38,160 | 17.20 |
| | | 2 | 99.16 | 37,293 | 17.20 |
| | | | 99.11 | 28,516 | 17.20 |
| | | | 99.10 | 38,987 | 17.20 |
| | | 1 | 98.91 | 38,377 | 17.20 |
| | | | 98.90 | 29,940 | 17.20 |
| | | | 98.44 | 36,986 | 17.30 |
| | | 0.2 | 96.76 | 39,740 | 17.10 |
| | | | 97.11 | 31,845 | 17.00 |

| | | | | | |
|---|---|---|---|---|---|
| | | | 95.84 | 36,444 | 17.50 |
| Contamination | NA12878 | 200 | 99.28 | 35,099 | 17.22 |
| | | 200 | 99.28 | 44,479 | 17.22 |
| | | 200 | 99.27 | 40,239 | 17.21 |
| | | 200 | 99.28 | 39,835 | 17.22 |
| | | 200 | 99.27 | 34,969 | 17.22 |
| | | 200 | 99.27 | 43,451 | 17.21 |
| | NA24385 | 200 | 99.92 | 37,549 | 16.85 |
| | | 200 | 99.92 | 42,925 | 16.84 |
| | | 200 | 99.91 | 42,820 | 16.85 |
| | NA24631 | 200 | 99.91 | 41,512 | 15.89 |
| | | 200 | 99.92 | 38,623 | 15.89 |
| | | 200 | 99.92 | 43,733 | 15.88 |
| | NC1 | UND | 64.15 | 767 | 42.89 |
| | NC2 | UND | 60.72 | 791 | 75.07 |
| | NC3 | UND | 60.43 | 784 | 69.69 |
| | NC4 | UND | 59.19 | 969 | 55.64 |
| | NC5 | UND | 63.36 | 1,127 | 40.19 |
| | NC6 | UND | 58.85 | 965 | 66.96 |
| | NC7 | UND | 59.28 | 930 | 43.62 |
| | NC8 | UND | 59.40 | 1,016 | 51.86 |
| | NC9 | UND | 58.80 | 872 | 71.14 |
| | NC10 | UND | 58.10 | 943 | 57.35 |
| | NC11 | UND | 61.35 | 1,129 | 70.60 |
| | NC12 | UND | 59.45 | 1,084 | 78.10 |
| | RB1 | UND | 60.61 | 829 | 75.95 |
| | RB2 | UND | 58.29 | 1,013 | 47.85 |
| | RB3 | UND | 60.11 | 1,156 | 55.88 |
| Degradation | NA12878 | 20 | 99.23 | 38,160 | 17.20 |
| | | | 99.24 | 39,176 | 17.20 |
| | NA12878 - 0 mJ/cm$^2$ | 0.2 | 97.07 | 40,268 | 17.00 |
| | | | 97.15 | 43,284 | 17.10 |
| | NA12878 - 125 mJ/cm$^2$ | 0.2 | 59.70 | 40,169 | 16.80 |
| | | | 59.56 | 42,016 | 17.60 |
| | NA12878 - 375 mJ/cm$^2$ | 0.2 | 50.33 | 32,739 | 10.60 |
| | | | 49.13 | 30,768 | 10.30 |
| | NA12878 - 625 mJ/cm$^2$ | 0.2 | 59.11 | 19,898 | 6.20 |
| | | | 62.60 | 20,202 | 6.40 |
| | NA12878 - 1000 mJ/cm$^2$ | 0.2 | 85.71 | 4,973 | 2.10 |
| | | | 83.99 | 3,917 | 2.80 |
| | NA24631 | 20 | 99.87 | 34,586 | 15.80 |
| | | | 99.85 | 36,783 | 15.80 |
| | NA24631 - 0 mJ/cm$^2$ | 0.2 | 97.69 | 41,107 | 15.70 |
| | | | 80.63 | 7,560 | 18.70 |
| | NA24631 - 125 mJ/cm$^2$ | 0.2 | 60.44 | 39,985 | 15.90 |
| | | | 59.68 | 40,184 | 16.40 |

| | | | | | |
|---|---|---|---|---|---|
| | NA24631 - 375 mJ/cm[2] | 0.2 | 49.76 | 33,089 | 10.50 |
| | | | 46.87 | 33,075 | 11.00 |
| | NA24631 - 625 mJ/cm[2] | 0.2 | 56.10 | 21,260 | 6.80 |
| | | | 55.22 | 21,188 | 6.80 |
| | NA24631 - 1000 mJ/cm[2] | 0.2 | 85.77 | 4,846 | 2.30 |
| | | | 86.63 | 37,666 | 3.00 |
| Species | Dog | 200 | 83.20 | 2,193 | 1.71 |
| | | | 79.85 | 2,035 | 1.59 |
| | Ecoli | 200 | 64.33 | 1,072 | 17.45 |
| | | | 65.54 | 749 | 92.19 |
| | Mouse | 200 | 74.08 | 1,718 | 12.29 |
| | | | 74.81 | 1,408 | 7.99 |
| | Rhesus | 200 | 62.60 | 34,080 | 5.77 |
| | | | 62.56 | 35,787 | 5.40 |
| | Yeast | 200 | 55.73 | 1,005 | 34.27 |
| | | | 63.49 | 776 | 90.41 |
| Mock Case | Mock1 | 3.66 | 88.38 | 37,298 | 18.67 |
| | Mock2 | 4.37 | 93.69 | 39,976 | 17.15 |
| | Mock5 | 0.85 | 68.93 | 42,793 | 18.24 |
| | Mock6 | 1.85 | 86.65 | 42,897 | 18.51 |
| | Mock7 | 9.61 | 83.16 | 44,983 | 19.39 |
| | Mock8 | 5.64 | 79.54 | 44,822 | 19.84 |
| | Mock9 | 62.92 | 89.95 | 46,836 | 17.83 |
| | Mock10 | 2.08 | 71.63 | 43,753 | 18.38 |
| | Mock13 | 0.36 | 0.00 | 25,645 | 16.85 |
| | Mock19 | 0.13 | 39.09 | 1,853 | 17.56 |
| | Mock20 | 0.13 | 68.19 | 1,824 | 24.51 |
| | Mock21 | 0.14 | 62.31 | 1,712 | 12.34 |
| Mixture | NA12878 | 200 | 99.29 | 31,598 | 17.21 |
| | | | 99.30 | 37,674 | 17.21 |
| | NA24631 | 200 | 99.94 | 37,720 | 15.89 |
| | | | 99.94 | 31,920 | 15.88 |
| | Mix1 (3:1) | 200 | 87.55 | 38,560 | 19.12 |
| | | | 87.66 | 34,031 | 19.11 |
| | Mix2 (9:1) | 200 | 97.20 | 39,584 | 16.21 |
| | | | 97.16 | 33,368 | 16.20 |
| | Mix3 (1:1) | 200 | 81.82 | 38,530 | 16.46 |
| | | | 82.36 | 34,906 | 17.08 |
| | Mix4 (1:3) | 200 | 90.25 | 27,586 | 18.85 |
| | | | 90.21 | 35,337 | 18.79 |
| | Mix5 (1:9) | 200 | 98.45 | 28,943 | 17.33 |
| | | | 98.50 | 37,292 | 17.32 |
| Reproducibility | NA12878 | 200 | 99.22 | 36,308 | 17.21 |
| | | 20 | 99.15 | 36,094 | 17.21 |
| | | 2 | 97.49 | 31,889 | 17.41 |
| | | 0.2 | 94.61 | 29,737 | 16.75 |

| | | | | | |
|---|---|---|---|---|---|
| | NA24385 | 200 | 99.90 | 36,431 | 16.85 |
| | NA24631 | 200 | 99.91 | 38,153 | 15.89 |
| | Mock1 | 3.66 | 95.98 | 31,623 | 18.55 |
| | Mock2 | 4.37 | 98.99 | 34,528 | 16.86 |
| | Mock6 | 1.85 | 85.31 | 29,199 | 18.17 |
| | Mock7 | 9.61 | 97.86 | 33,210 | 18.02 |
| | Mock8 | 5.64 | 94.60 | 34,615 | 17.89 |
| | Mock9 | 62.92 | 99.84 | 35,026 | 16.86 |
| | Mock10 | 2.08 | 85.94 | 35,813 | 17.06 |

<u>Table S3</u>

Samples with heterozygosity outside the 15-20% range, including negative controls (NCs), reagent blanks (RBs), non-human DNA, and highly degraded or low quantity human samples.

| Study | Sample ID | DNA Input into GSA (ng) | Call Rate (%) | Total Intensity | Heterozygosity (%) |
|---|---|---|---|---|---|
| Degradation | NA12878 - 375 mJ/cm$^2$ | 0.2 | 50.33 | 32,739 | 10.60 |
| | | | 49.13 | 30,768 | 10.30 |
| | NA12878 - 625 mJ/cm$^2$ | 0.2 | 59.11 | 19,898 | 6.20 |
| | | | 62.60 | 20,202 | 6.40 |
| | NA12878 - 1000 mJ/cm$^2$ | 0.2 | 85.71 | 4,973 | 2.10 |
| | | | 83.99 | 3,917 | 2.80 |
| | NA24631 - 375 mJ/cm$^2$ | 0.2 | 49.76 | 33,089 | 10.50 |
| | | | 46.87 | 33,075 | 11.00 |
| | NA24631 - 625 mJ/cm$^2$ | 0.2 | 56.10 | 21,260 | 6.80 |
| | | | 55.22 | 21,188 | 6.80 |
| | NA24631 - 1000 mJ/cm$^2$ | 0.2 | 85.77 | 4,846 | 2.30 |
| | | | 86.63 | 37,666 | 3.00 |
| Contamination | NC1 | UND | 64.15 | 767 | 42.89 |
| | NC2 | UND | 60.72 | 791 | 75.07 |
| | NC3 | UND | 60.43 | 784 | 69.69 |
| | NC4 | UND | 59.19 | 969 | 55.64 |
| | NC5 | UND | 63.36 | 1,127 | 40.19 |
| | NC6 | UND | 58.85 | 965 | 66.96 |
| | NC7 | UND | 59.28 | 930 | 43.62 |
| | NC8 | UND | 59.40 | 1,016 | 51.86 |
| | NC9 | UND | 58.80 | 872 | 71.14 |
| | NC10 | UND | 58.10 | 943 | 57.35 |
| | NC11 | UND | 61.35 | 1,129 | 70.60 |
| | NC12 | UND | 59.45 | 1,084 | 78.10 |
| | RB1 | UND | 60.61 | 829 | 75.95 |
| | RB2 | UND | 58.29 | 1,013 | 47.85 |
| | RB3 | UND | 60.11 | 1,156 | 55.88 |
| Species | Dog | 200 | 83.20 | 2,193 | 1.71 |
| | | 200 | 79.85 | 2,035 | 1.59 |
| | Ecoli | 200 | 65.54 | 749 | 92.19 |

| | | | | | |
|---|---|---|---|---|---|
| | Mouse | 200 | 74.08 | 1,718 | 12.29 |
| | | 200 | 74.81 | 1,408 | 7.99 |
| | Rhesus | 200 | 62.60 | 34,080 | 5.77 |
| | | 200 | 62.56 | 35,787 | 5.40 |
| | Yeast | 200 | 55.73 | 1,005 | 34.27 |
| | | 200 | 63.49 | 776 | 90.41 |
| Mock Case | Mock20 | 0.13 | 68.19 | 1,824 | 24.51 |
| | Mock21 | 0.14 | 62.31 | 1,712 | 12.34 |

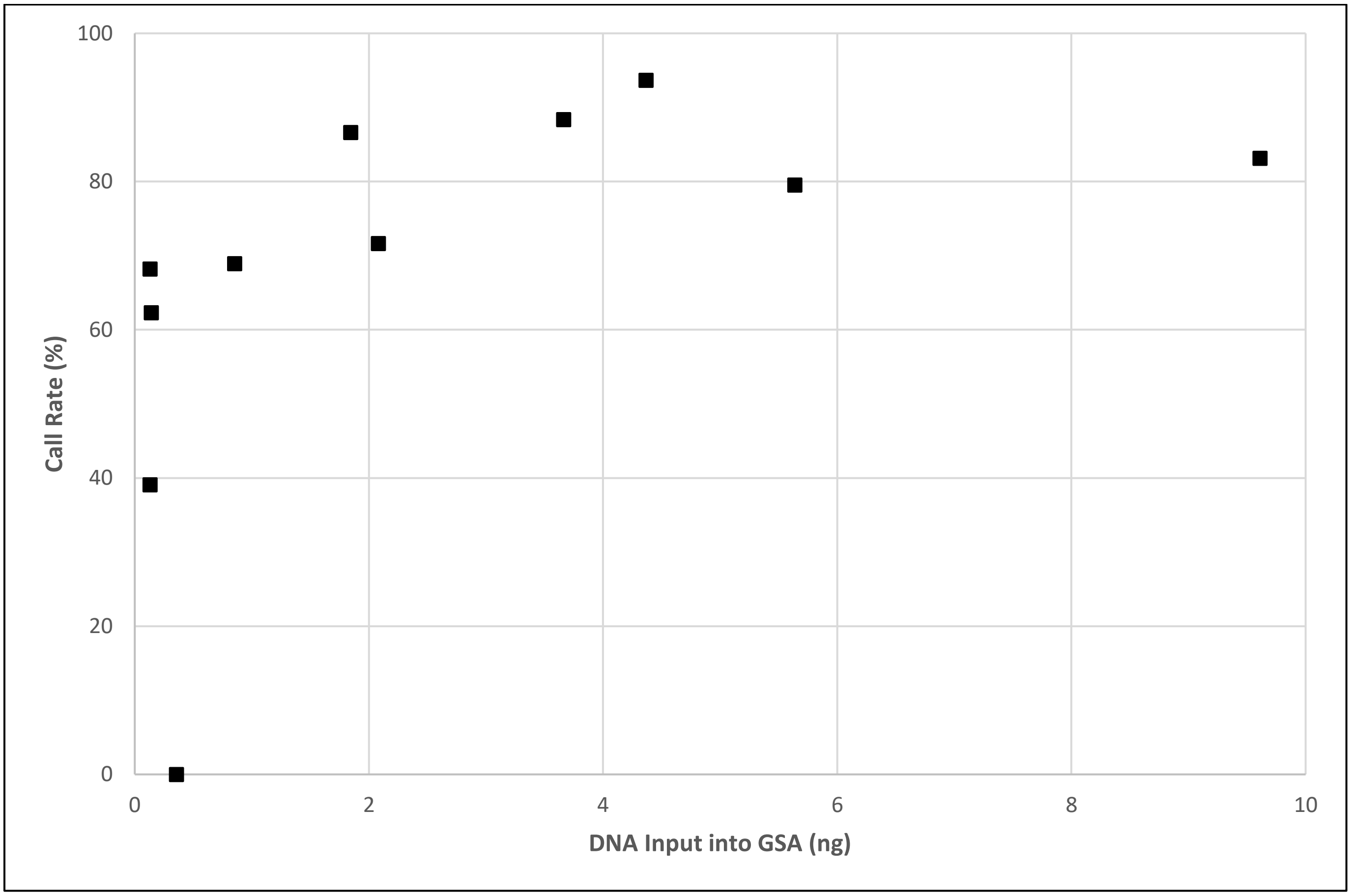


<u>Figure S2</u>
Call rates generated from mock samples with various DNA input. Sample Mock9 was an outlier with nearly 63 ng of DNA input and was excluded from this figure.

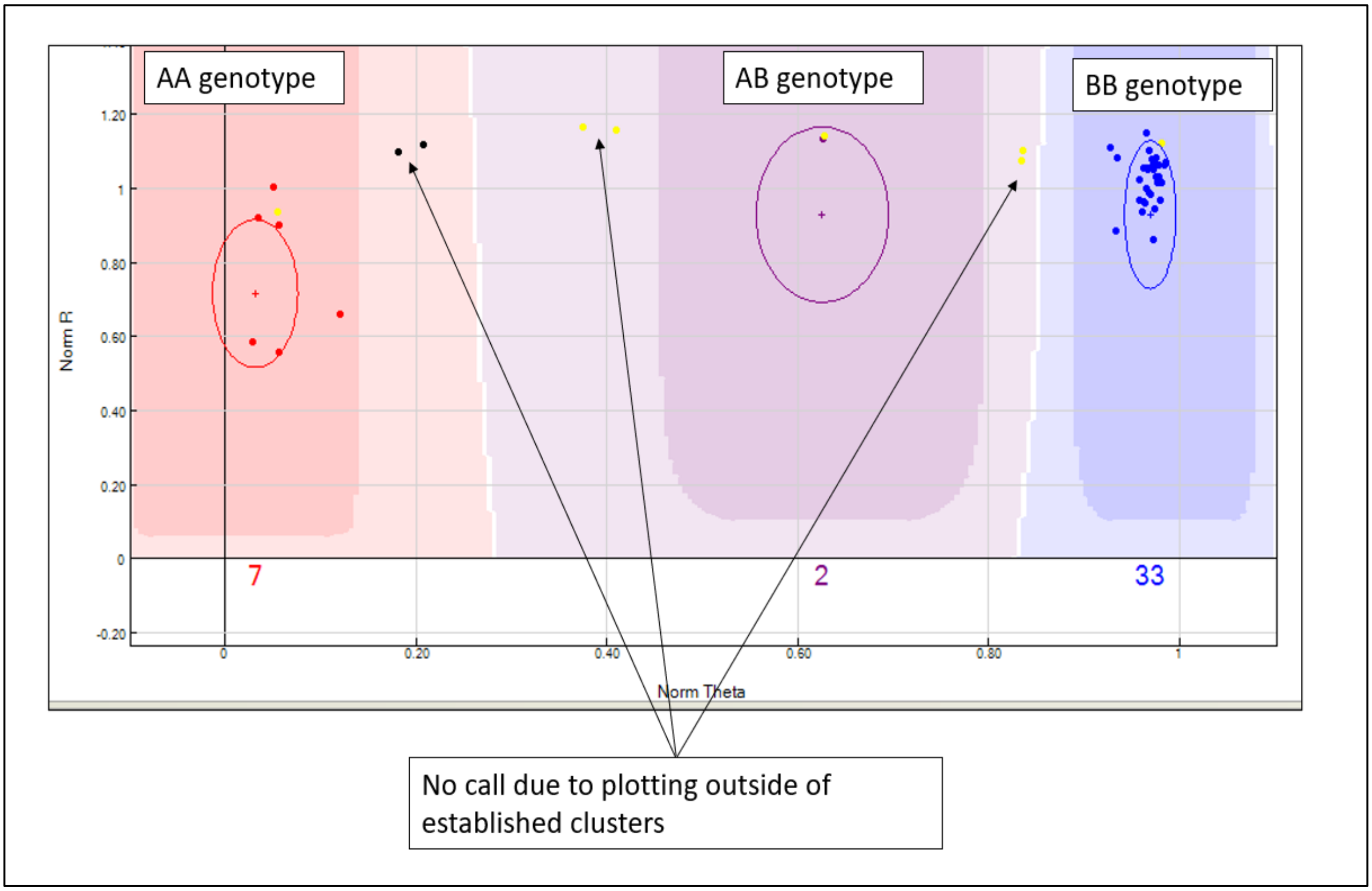


<u>Figure S3</u>

Example of established clusters in GenomeStudio for genotypes AA, AB, and BB. Calls that do not fall neatly into one of the expected clusters (dark shaded regions) results in a no call at that locus. Each dot represents a single sample at a particular locus.

**Table S4**

Sample concordance between replicates processed as part of the repeatability and reproducibility studies.

| Study | Sample ID | Concordance (%) |
|---|---|---|
| Reproducibility | Mock1 | 98.69 |
| | Mock2 | 99.14 |
| | Mock6 | 99.03 |
| | Mock7 | 96.06 |
| | Mock8 | 95.88 |
| | Mock9 | 97.70 |
| | Mock10 | 94.28 |
| | NA12878 - 200 ng | 100.00 |
| | NA12878 - 20.0 ng | 100.00 |
| | NA12878 - 2.0 ng | 99.81 |
| | NA12878 - 0.2 ng | 99.64 |
| | NA24385 - 200 ng | 100.00 |
| | NA24631 - 200 ng | 100.00 |
| Repeatability | NA12878 - 200 ng | 100.00 |
| | | 100.00 |
| | | 100.00 |
| | | >99.99 |
| | | >99.99 |
| | NA24385 - 200 ng | >99.99 |
| | | >99.99 |
| | NA24631 - 200 ng | >99.99 |
| | | 100.00 |
| | NA12878 - 20.0 ng | >99.99 |
| | | >99.99 |

**Table S5**

Re-scan information for BeadChips and samples included in the stability study.

| BeadChip | Study | Time Between Scans | Number of Samples |
|---|---|---|---|
| 1 | Reproducibility | 1 day | 22 |
| | Reproducibility | 1 week | 22 |
| 2 | Reproducibility | 1 month | 23 |
| 3 | Species | 3 months | 12 |
| 4 | Degradation | 1 year | 24 |
| 5 | Sensitivity | 2 years | 21 |

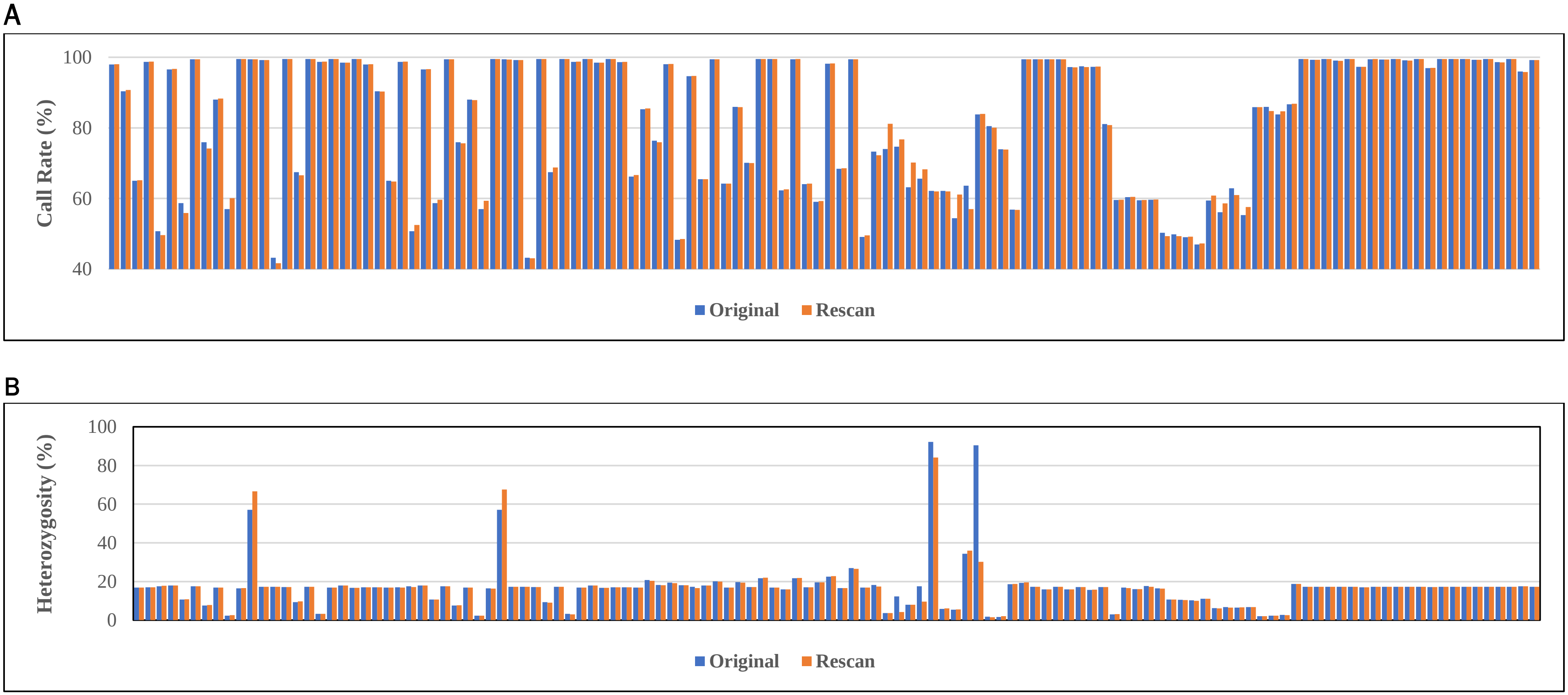


Figure S4

Call rate (A) and heterozygosity (B) generated from the original scan and re-scan data for 124 samples used in this study.
